# Differences between human male and female neutrophils with respect to which RNAs are present in polysomes

**DOI:** 10.1101/2025.05.07.652701

**Authors:** Darrell Pilling, Kristen M. Consalvo, Sara A. Kirolos, Richard H. Gomer

## Abstract

**Background:** Human males and females show differences in the incidence of neutrophil-associated diseases such as systemic lupus erythematosus, rheumatoid arthritis, and multiple sclerosis, and differences in neutrophil physiological responses such as responses to infection, tissue damage, and chemotactic factors. However, little is known about the basis of sex-based differences in human neutrophils.

**Methods:** Starting with human neutrophils from healthy donors, we used RNA-seq to examine total RNA profiles, RNAs not associated with ribosomes and thus not being translated, RNAs in monosomes, and RNAs in polysomes and thus heavily translated.

**Results:** There were sex-based differences in the levels of RNAs across free RNA, monosome, and polysome fractions. Male neutrophils had increased levels of mRNAs encoding mitochondrial proteins in the free RNA fractions, indicating low levels of translation. The polysomes of male neutrophils were enriched for mRNAs encoding cytoskeletal organization, cell motility, and cell activation. The polysomes of female neutrophils were enriched for mRNAs associated the regulation of metabolic processes, cytokine responses, and mitochondrial proteins.

**Conclusions:** These data indicate that male and female neutrophils have different expression patterns and different translation efficiency of some mRNAs. This may contribute to the observed sex-based differences in neutrophil behavior and neutrophil-associated disease incidence and severity.

## Introduction

The mammalian immune system displays widespread sexual dimorphism ^1, 2^. Females tend to have more robust immune responses than males ^2–4^, and reduced rates of infections ^5, 6^. Females have a higher incidence of autoimmune disorders compared to males, and an increase in post-transplantation organ rejection ^4, 7, 8^. Some of these sex differences can be explained by hormonal differences ^9^ or sex chromosome copy number ^10^, but there is much that is still unknown ^4^.

Neutrophils are the most abundant circulating immune cell in humans, representing 50-70% of all leukocytes and are a major component of the innate immune system ^11, 12^. Neutrophils also have a role in tissue homeostasis, but aberrant activation and persistence can contribute to inflammation and the progression of some diseases^11, 12^ including acute respiratory distress syndrome (ARDS), rheumatoid arthritis (RA), and other disorders ^11–14^.

In human circulating neutrophils, there are sex-based differences in phenotype and function, with adult female neutrophils having a more mature phenotype as determined by surface receptor expression, enhanced type I interferon pathway activity, and increased response to GM-CSF compared to adult male neutrophils ^15–18^. There are also sex-based differences in the levels of some proteins, with females having higher levels of proteins that regulate metabolic processes and proteolytic pathways in their neutrophils, and males having more translational control and signaling proteins in their neutrophils ^19^. Males have more phosphorylation of proteins that inhibit transcription, regulate protein localization and apoptotic signaling in mitochondria, and modulate neutrophil activation ^19^. Although female neutrophils expressed many phosphoproteins, there were no phosphoproteins that were significantly changed compared to male neutrophils ^19^. How these differences between male and female neutrophils is influenced by gene expression and translation is unclear.

To assess translation efficiency in circulating human neutrophils, we isolated neutrophils from male and female donors, lysed the cells, and separated the lysates on sucrose gradients to fractionate free RNA, RNA associated with monosomes, and RNA associates with polysomes. In this report, we describe, for human neutrophils, sex-based differences in gene expression and translation efficiency. These differences may contribute to the observed sex-based differences in neutrophil-associated disease incidence and severity.

## Methods

### Neutrophil isolation

Human venous blood was collected with the approval from the Texas A&M University Institutional Review Board (IRB number 2017-0792D) from healthy volunteers who gave written consent. Neutrophils were isolated as previously described ^20–22^. Briefly, neutrophils were isolated at room temperature (RT) from blood collected directly into vacutainer tubes containing EDTA (454209; Greiner Bio-One; Monroe, NC). Neutrophils were isolated by Polymorphprep (1114683; Axis-Shield, Oslo, Norway) following the manufacturers’ instructions, except the centrifugation of gradients was done for 40 minutes. After centrifugation, first the upper mononuclear band was removed and then a sterile wide-bore plastic pipette (414004-005, VWR, Radnor, PA) was used to collect the lower neutrophil band. Neutrophils were then mixed with 10 mL PBS (17-5 F; Lonza, Walkersville, MD) and collected by centrifugation for 10 minutes at 300 x g at RT. Neutrophil cell pellets were resuspended in PBS and then recentrifuged. The resuspension and centrifugation was repeated 3 more times.

### RNA and ribosome collection, fractionation, and purification

From each donor, pellets of 45 to 115 x 10^6^ neutrophils were disrupted by pipetting vigorously with 500 µl ice cold “Complete Polysome Buffer” (15 mM Tris-HCl pH 7.5, 300 mM NaCl, 15 mM MgCl_2_, 1% Triton X-100 (Alfa Aesar, Ward Hill, MA), 100 µg/ml Cycloheximide (VWR, Radnor, PA), 1 mg/ml Heparin (A16198.06, Thermo Scientific, Rockford, IL), 500 units/ml RNasin Ribonuclease inhibitor (Invitrogen, Carlsbad, CA), 20 mM DTT, and 10x Protease and Phosphatase inhibitor cocktail (Thermo Scientific)) ^23^. Lysed samples were separated on a 10-50% sucrose gradient made with Polysome Gradient Buffer (10 mM HEPES-KOH pH 7.5, 70 mM ammonium acetate, 5mM magnesium acetate, prepared the same day) as previously described ^23^. Cell lysates were layered on top of the prepared sucrose gradient, centrifuged, and then fractionated following the manufacturer’s instructions for a TriAX flow cell (BioComp, Fredericton, New Brunswick, Canada) and FC203B fraction collector (Gilson, Middleton, WI). Fractions were collected and combined so that F1-3 contained free RNA, F4-6 contained monosome associated RNA, F7-9 contained early polysome associated RNA, and F10-12 contained late polysome associated RNA. RNA purification and precipitation was performed as described ^23^. Briefly, 0.5 ml of each sucrose fraction was mixed with 0.5 ml TRIzol (Invitrogen) and 0.2 ml chloroform, then clarified by centrifugation at 12,000 x g for 15 minutes at 4°C. 0.5 ml of the upper layer was transferred to a fresh tube containing 1 ml isopropanol and 2 µl of 15 mg/ml Glycoblue (Invitrogen). After mixing, the RNA was precipitated by incubating overnight at -20°C and collected by centrifugation at 12,000 x g for 15 minutes at 4°C. The pellet was rinsed with 1 ml ice-cold 70% ethanol. The ethanol was removed after centrifugation at 12,000 x g for 15 minutes at 4°C. Precipitated samples were re-spun a second time to remove the remaining ethanol from the side of the sample tubes. RNA pellets were air dried for at least 10 minutes at room temperature before being dissolved in 20 µl nuclease-free water (Thermo Scientific). RNA concentrations were checked with a Synergy Mx plate reader with a microdrop attachment (BioTek, Winooski, VT).

### RNA sequencing

RNAseq libraries were created following the manufacturer’s instructions for QuantSeq 3’ mRNA-Seq Library Prep Kit FWD for Illumina (type 015.96, Lexogen Inc, Greenland, NH), with 2 µg of RNA used as the starting material. Libraries were sequenced using an Illumina NextSeq 500 platform (Texas A&M University Institute for Genome Sciences and Society Experimental Genomics Core, College Station, TX). RNA sequencing data were analyzed using the QuantSeq Data Analysis Pipeline on the BlueBee Genomic Platform (BlueBee, San Mateo, CA). The quality of sequences was evaluated using FastQC software (version 0.11.5) after adapter trimming with BBDUK software (version 35.92). Gene and transcript intensities were computed using STAR software (version 2.5.2a) with the Gencode Release 27 (GRCh38) human genome as a reference. Raw files of RNA-Seq data were uploaded to GEO (accession number GSE288976).

### Data analysis

For each donor, for each specific RNA (indicated by X), the normalized count of X in the free fraction was calculated as (read count of X in the free fraction)/ (total number of read counts in the free fraction). The normalized count of X in the monosome fractions was similarly calculated as (read count of X in the monosome fraction)/ (total number of read counts in the monosome fraction). Similar normalization was done for early polysomes and late polysomes. For the free/ monosome/ polysome analysis, there were 4 female and 3 male donors. We only analyzed RNAs where the RNA was detected in one fraction (F1-3, F4-6, F7-9, or F10-12) for at least 3 of the 4 females, or all 3 of the males. However, most of the RNAs analyzed were present in multiple fractions.

Differences in RNA expression between free RNA, and RNA associated with monosomes, and early and late polysome in males and females was assessed using t-tests. Fold change in expression and t-test values were ranked for volcano plot visualization. Gene ontology (GO) pathway analysis was performed, and graphs were generated, using ShinyGO (v 0.82 using Ensembl Release 107) ^24^. Groups were analyzed with the standard “Use pathway DB for gene counts” in the Homo sapiens database, and significance (p < 0.05) was determined by Fisher’s exact test with FDR correction. Results were confirmed using g:Profiler (https://biit.cs.ut.ee/gprofiler/gost) and Metascape (https://metascape.org/). Venn diagrams were generated using InteractiVenn (www.interactivenn.net). Principal Component Analysis (PCA) and heatmaps were generated using ClustVis (https://biit.cs.ut.ee/clustvis_large/) ^25^.

### Statistics

Prism v7 (GraphPad Software Inc., San Diego, CA, USA) and Microsoft 365 Excel (Microsoft, Redmond, WA) were used for data analysis. Graphs were generated with Prism. Data are shown as mean ± SEM except where otherwise stated. To determine whether the mean difference between two groups was statistically significant, the Mann-Whitney test was used. Statistical significance was defined as p 0.05. For the volcano plots, one unpaired t test per row was calculated, without assuming consistent SD (the fewer assumptions option), with an uncorrected significance of p < 0.05 considered significant.

## Results

### As previously observed, male and female neutrophils show differences in levels of some RNAs

The transcriptome of human and murine neutrophils has been relatively well studied ^26–29^. Previous studies also found sex-based differences in human neutrophils ^15, 18, 30^. To confirm this, we analyzed freshly isolated human neutrophils from 2 healthy male and 4 female donors by RNA-seq and identified 2,175 transcripts from datasets that had detectable RNA sequences in either both male donors or in at least 2 of 4 female donors **(Table S1 Tab 1)** ^31^. There was no significant difference in the number of detected RNA transcripts between the male and female samples (**Figure S1A**) ^31^. As described below in the results sections “Some transcripts were only present in male neutrophils”, and “Some transcripts were only present in female neutrophils”, we observed some RNAs in this data set that appeared to be present only in male or only in female neutrophils (**Table S1 Tabs3 and 4)** ^31^. In addition to RNAs present only in males or only in females, there were 53 transcripts with greater abundance in male cells and 1 transcript (RAB21) with greater abundance in female cells (**Figure 1A and Table S1 Tab5**) ^31^. GO term pathway analysis of the proteins encoded by the 53 male transcripts were enriched for mRNAs encoding proteins involved with regulation of translation and protein targeting to the endoplasmic reticulum (RPL5, RPL35, RPLP1, RPS3, and DDX3X), lamellipodia and focal adhesion (PABPC1, PFN1, RPL5, CAP1, RDX, RPLP1, ACTR2, RPS3, SLA, and JAK1), and secretory granules (CAP1, ACTR2, SRP14, ANXA2, and DDX3X) (**Figure 1B**). These results are consistent with previous data indicating that male and female neutrophils have transcriptomic differences ^15, 18, 30^.

**Figure 1.**
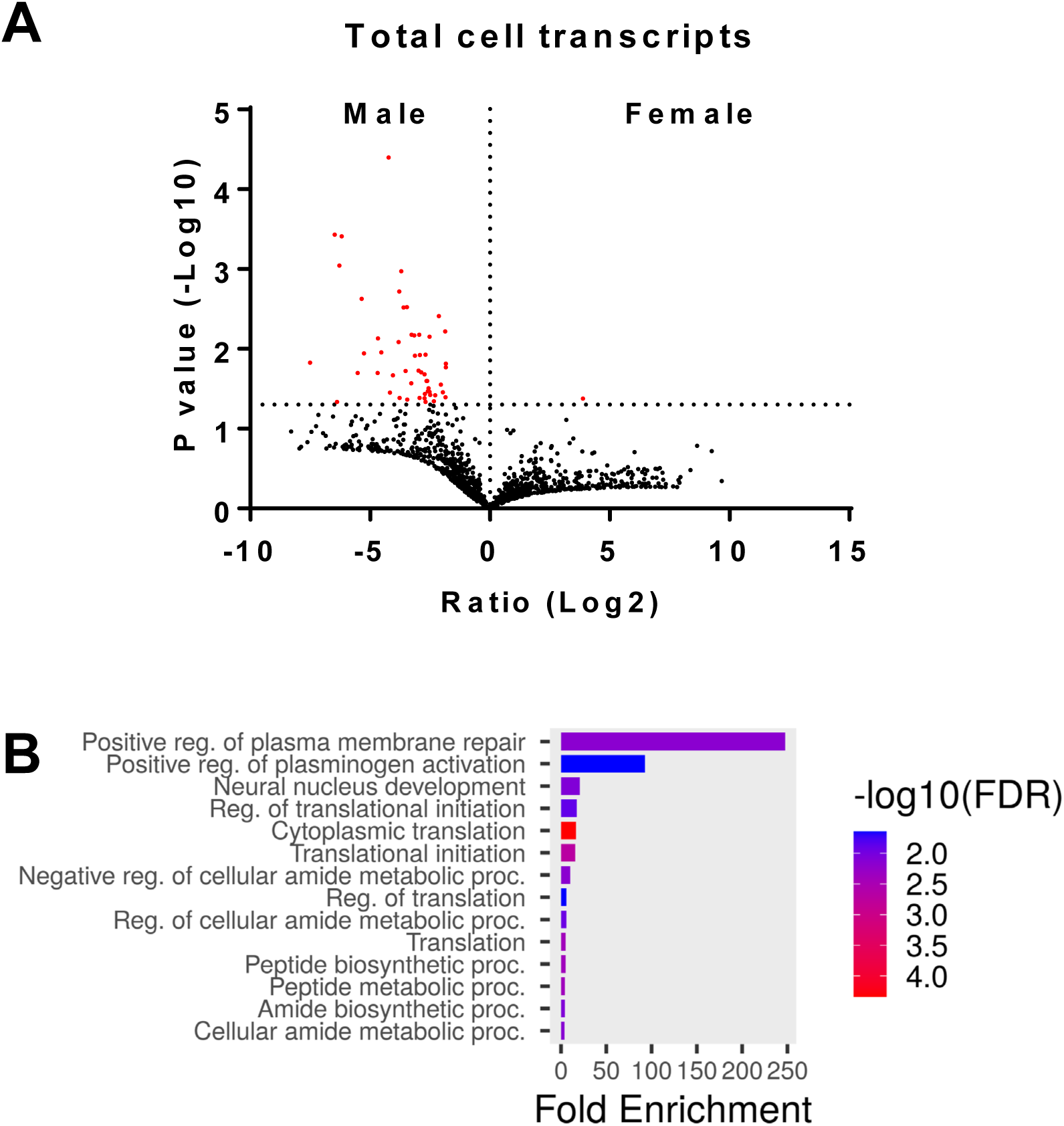
Male and female neutrophils have differences in the total levels of some RNAs. **A)** Volcano plot showing the fold change (Log2) and p-value (-Log10) comparing the total transcriptomes from male and female donors. Transcripts are marked in red have p values <0.05 (-Log10 >1.3). **B)** Gene ontology analysis indicates that some mRNAs present at higher levels in male neutrophils encode proteins associated with translation regulation.

### Polysome fractionation of male and female neutrophils

Polysome fractionation and profiling has been used to analyze translation efficiency in many cell types, including human platelets, monocyte cell lines, and macrophages ^32–36^. However, RNA-seq analysis after polysome fractionation is rare in neutrophils with the human promyelocytic leukemia HL-60 cell line being used as a surrogate for neutrophils ^33, 37^. To assess translation efficiency in circulating human neutrophils, we isolated neutrophils from 3 healthy male and 4 healthy female donors, lysed the cells, and separated the lysates on sucrose gradients ^23, 38^. Fractionated male and female neutrophils, despite showing donor to donor variations in the profiles, all contained a clear monosome peak (**Figure 2**). Similar experiments on the human MCF7 cancer cell line also showed replicate experimental variation in the ribosome profiles ^39^. The coefficient of variation (Standard Deviation / Mean) for the polysome region (defined as gradient position 40 – 75, consisting of fractions 7 - 12), showed no significant difference between the male and female profiles. These profiles show some indication of peaks in the polysome regions for both male and female neutrophils, most clearly seen in male donor #2 and female donors #1 and #4 (**Figure 2**). Neutrophils have a significantly lower resting gene expression profile than other immune cell types, such as peripheral blood mononuclear cells ^40^, with low but detectable transcriptional activity ^41, 42^, which increases rapidly after neutrophil activation ^41^. This reduced basal transcription activity may be responsible for the relatively low polysome peaks.

**Figure 2.**
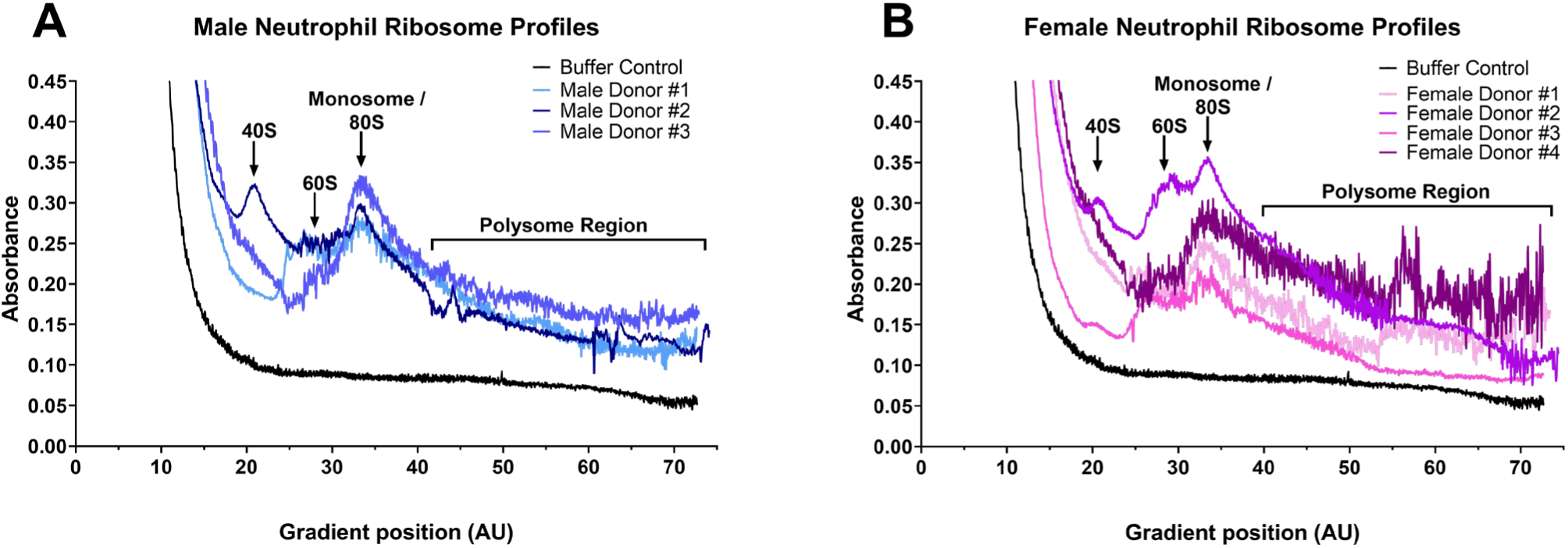
Free, monosome, and polysome fractionation. Male and female neutrophils were isolated from blood, lysed, the contents were separated on a sucrose gradient, and the gradient was then fractionated. Results are separated into **(A)** male donors (n = 3) and **(B)** female donors (n = 4).

### Male and female neutrophils show differences in RNA transcripts across polysome fractions

We then analyzed RNA transcripts from the polysome fractions. RNA-seq data was collated for free RNA (Fractions 1 – 3, corresponding to gradient positions 1 – 20), monosomes (Fractions 4 – 6, corresponding to gradient positions 21 – 40), early polysomes (Fractions 7 – 9, corresponding to gradient positions 41 – 55), and late polysomes (Fractions 10 – 12, corresponding to gradient positions 56 – 75) (**Figures 2A and B**). Analysis was restricted to datasets where we identified transcripts from all 3 male donors or at least 3 of the four female donors in at least one fraction. We identified 3,056 transcripts from datasets that had transcripts in at least one fraction (F1-3, F4-6, F7-9, or F10-12) (**Table S2 Tab 1)** ^31^. There were no significant differences in the total number of detected RNA transcripts in the combined fractions between the male and females (**Figure S1B**) ^31^, but there were more transcripts in the monosome fraction (F4-6) compared to the other 3 fractions (**Figures S1C**) ^31^.

Principal-component analysis (PCA) highlighted clear segregation of RNA into free, monosome, early polysome, and late polysome fractions, and a general trend that male and female free RNA and monosome fractions were more closely associated, but the male and female early and late polysomes were more diverse (**Figure 3A**). Using unsupervised clustering (heatmap), free RNA and monosome fractions were clustered together, with male and female early and late polysome fractions being more diverse, with early polysome fractions having the greatest number of diverse transcripts (**Figure 3B**).

**Figure 3.**
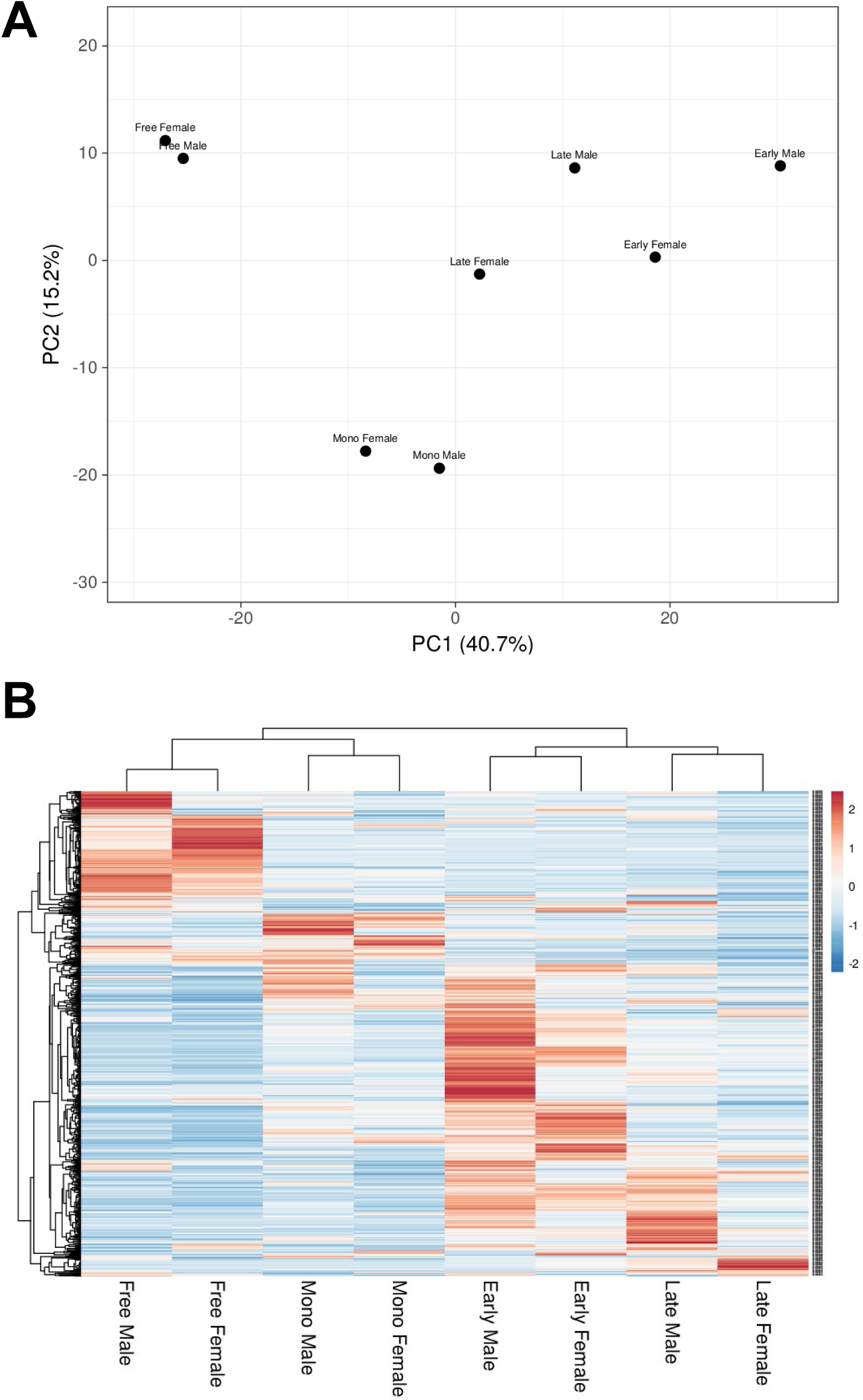
Analysis of RNA transcriptomes from free RNA, monosomes, and polysomes of male and female donors. **A)** Principal component analysis based on per-sample mean-centered log2 values of male and female transcripts. **B)** Heat map showing normalized expression levels (z score) of RNAs in fractions from male and female neutrophils.

### Male/female differences in RNAs in the Free RNA fraction

In the free RNA fraction (F1-3), there were 41 RNAs with significantly greater abundance in males, and 6 RNAs with greater abundance in females (**Figure 4A and Table S2 Tab2**) ^31^. In the male free RNA fraction, which represent transcripts that are not being translated into protein, there were 40 transcripts with levels > 2 fold higher than the average transcript level (Z>2). These 40 transcripts were enriched for mRNAs encoding mitochondrial respiratory chain complex proteins (APP, CEBPA, COX5B, COX7A2, LDHA, MT-CO2, and NCF2) (**Figure 5A and Table S2 Tab 5**) ^31^. There were 45 Z>2 transcripts in the female free RNA fraction. These included mRNAs encoding proteins associated with RNA processing and regulation including AGO4, DDX3X, HNRNPA2B1, PKM, RPS3, RPL38, and RACK1) (**Figure 5B and Table S2 Tab 5**) ^31^.

**Figure 4.**
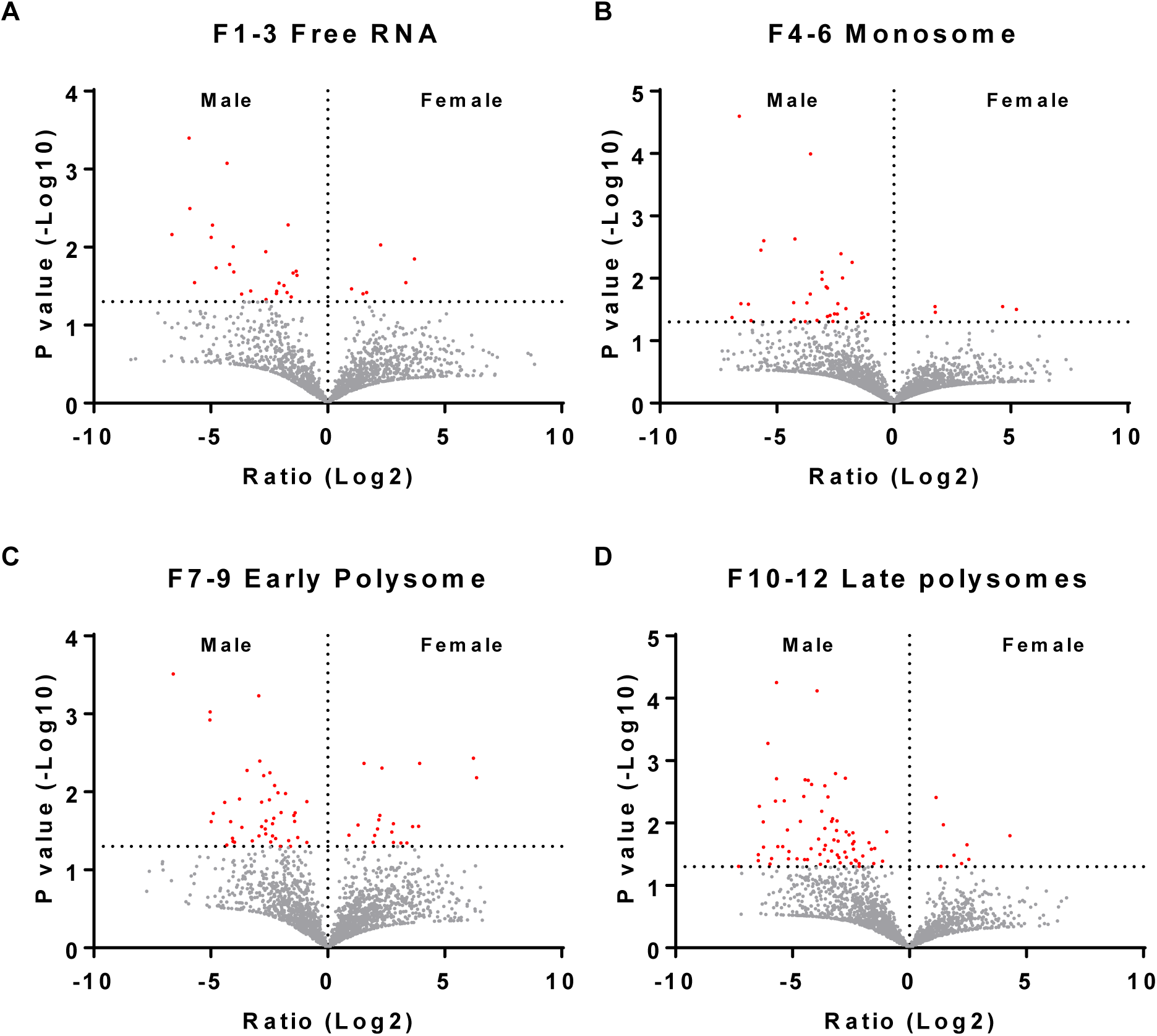
Comparison of transcriptomes from free RNA, monosomes, and early and late polysomes of male and female donors. Fractionated samples were combined into “Free RNA”, “Monosome”, “Early Polysome”, and “Late Polysome” samples for each donor. The samples were then treated to isolate, purify, and sequence mRNAs. Volcano plots from **A)** free RNA, **B)** monosomes, **C)** early polysomes and **D)** late polysome datasets showing the fold change (Log2) and p-value (-Log10) comparing the transcriptomes from male and female donors. RNA transcripts marked in red have p values <0.05 (-Log10 >1.3).

**Figure 5.**
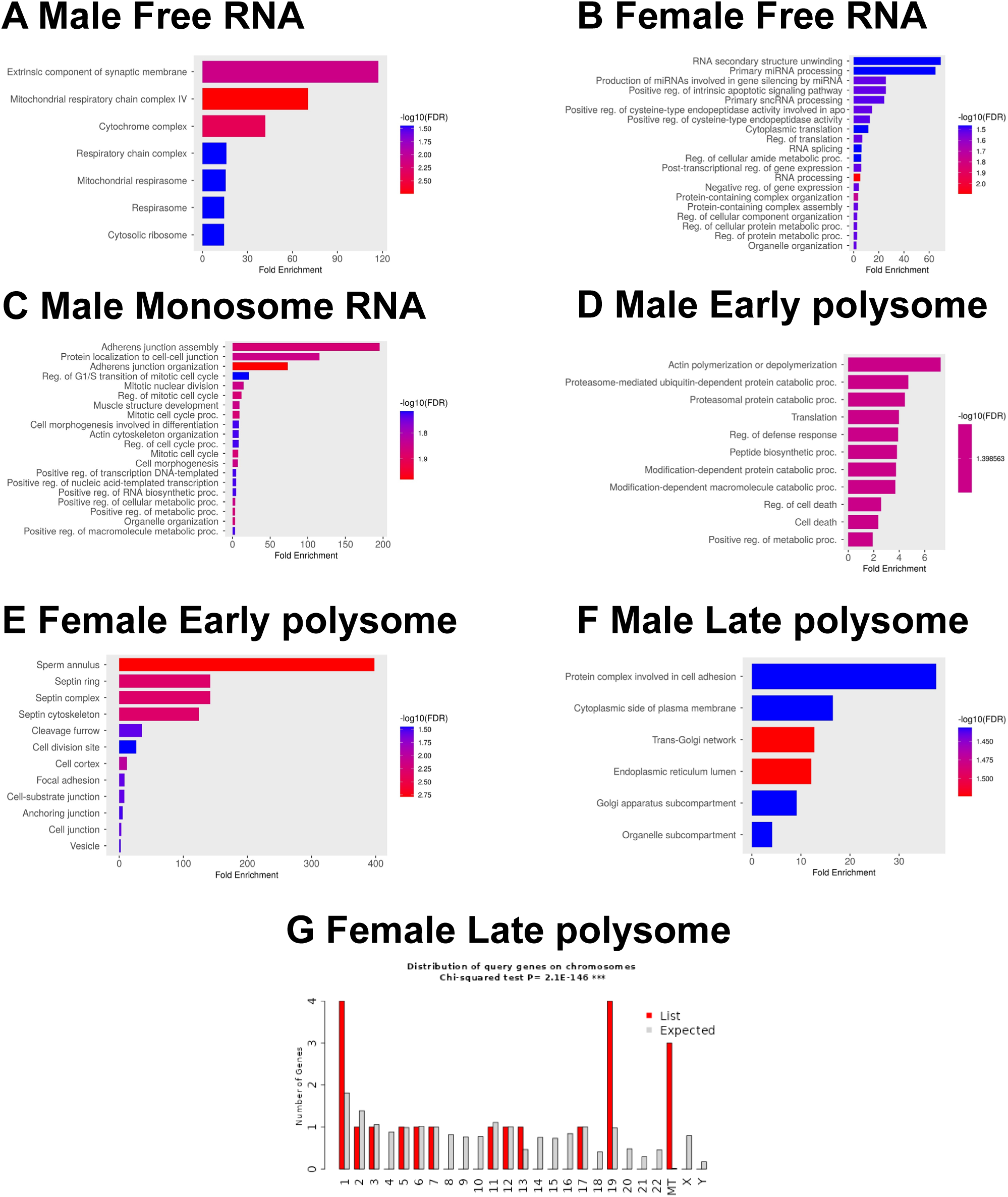
Analysis of heatmap data of transcripts with >2 fold change in male and female neutrophils. GO term analysis of enriched transcripts in free RNA from **A)** male and **B)** female neutrophils. **C)** Enriched transcripts in monosomes from males. Enriched transcripts in early polysomes from **D)** males and **E)** females. **F)** Enriched transcripts in late polysome fractions from males. **G)** Chi-squared analysis of the distribution by chromosome of enriched transcripts in late polysome fractions from females.

### Male/female differences in RNAs in the Monosome fraction

There were 44 RNAs with significantly greater abundance in the monosome fraction in males, and four with greater abundance in females (**Figure 4B and Table S2 Tab2**) ^31^. In the male monosome fraction, there were 20 Z>2 transcripts, including transcripts related to adhesion, cytoskeletal organization, and cell-cell interaction (including ACTB, ACTN4, ARHGAP25, CDC42, and PAK2) (**Figure 5C and Table S2 Tab 5**) ^31^. There were only 8 Z>2 female monosome transcripts, with no discernable enrichment (**Table S2 Tab 5**) ^31^.

### Male/female differences in RNAs in the Early Polysome fraction

There were 56 RNAs with greater abundance in the early polysome fraction in males, and 19 with greater abundance in females (**Figure 4C and Table S2 Tab2**) ^31^. There were 90 Z>2 male early polysome transcripts, and these were enriched for actin polymerization (COTL1, MTPN, DIAPH1, CFL1, RASA1, and ARPC3, catabolic processes (HSPA5 UBE4A, FAF2, FBXL5, TRIP12, PSMD1, CSNK1A1, UBB, ARRB2, and PCBP2), RNA processing and regulation (RPLP0, EEF1D, TARS1, CPEB4, RPL30, RPS21, UBA52, TPR, RPUSD4, CELF1, MTPN, and RRBP1), and vesicle regulation (HSPA5, SPEN, SLC25A3, HSP90AA1, RPLP0, ST13, COTL1, ARPC3, TARS1, ATP6V1B2, RPL30, MNDA, RAB8B, UBB, CFL1, RHOG,

P4HB, PCBP2, and UBA52) (**Figure 5D and Table S2 Tab 5**) ^31^. There were 24 Z>2 female early polysome transcripts, including transcripts related to septin protein complexes (SEPTIN6 and SEPTIN7) and adhesion (RPS5, ANXA5, PPP1CB, PTPRC, SEPTIN6, MTDH, PRKCB, and CACNA1E) (**Figure 5E and Table S2 Tab 5**) ^31^.

### Male/female differences in RNAs in the Late Polysome fraction

There were 109 RNAs with greater abundance in the late polysome fraction in males, and 8 with greater abundance in females (**Figure 4D and Table S2 Tab2**) ^31^. There were 24 Z>2 male late polysome transcripts which likely represent transcripts that are highly likely to be translated into protein, including transcripts related to cell adhesion (ITGA6 and LGALS1) and surface receptors (FCGR3B), the Golgi and ER network (TGOLN2, SORL1, PLPP3, ATXN2, KTN1, CKAP4, and COPB2), and RNA binding (LGALS1, PRRC2C, KTN1, SRRM1, CKAP4, ATXN2, POLE3, NACC2, SUZ12, DDX60L, KAT6A, and ETV5) (**Figure 5F and Table S2 Tab 5**) ^31^. Females had 20 Z>2 late polysome transcripts, and these were highly enriched for proteins and RNAs associated with mitochondria (MT-CYB, MT-RNR2, and MT-TH) (**Table S2 Tab 5 and Figure 5G**) ^31^.

### Male and female neutrophils show differences in enrichment of RNA transcripts in polysomes

To further elucidate sex-based differences in the translation of neutrophil RNAs, for each RNA from each donor (using only RNAs where the RNA was detected in one fraction (free, monosome, early polysome, or late polysome) for at least 3 of the 4 females, or all 3 of the males) we assessed the Total Translation Rate (TTRx) using: TTRx= (Monosome + Early Polysome + Late Polysome) / Free RNA.

For RNAs where all donors had a non-infinite value for TTRx, the mean and standard deviation was calculated for the TTRx value for each RNA (**Table S3 Tabs 1 and 2**). We identified 29 transcripts that were significantly different between male and female neutrophils using these criteria, with 20 RNAs (all mRNAs) having a higher TTRx in males and 9 higher in females (**Table S3 Tabs 2 and 3**) ^31^. The 20 mRNAs in males encoded proteins involved with vesicles including major vault protein (MVP), RAS family members (RAB27A and RIN3), a variety of enzymes (ALDOA and CANT1) and signaling molecules (DNM2, OSBPL2, and PEBP1) along with transcription factor complex nuclear factor-kappa-B (NFkB2), Lymphotoxin beta (LTB), and S100A2. The 9 female RNAs were enriched for interferon gamma signaling (SP100 and NMI), eukaryotic translation (EIF3A), and nuclear proteins (ARID4B, SP100, EIF3A, NMI, TKT, TOP3A, and ANP32A) (**Table S3 Tab 3**) ^31^.

We also assessed for each RNA X of each donor a Moderate Translation Rate (MTRx), (**Table S3 Tab 4**) ^31^, using:

MTRx = (Early Polysome + Late Polysome)/ (Free RNA + Monosome). We identified 49 RNAs that were significantly different between male and female neutrophils using these criteria (**Table S3 Tabs 5 and 6**) ^31^. Of these 49 RNAs, 37 were significantly upregulated in male samples and 12 were upregulated in female samples. The 37 upregulated transcripts in males were all mRNAs and were enriched for cell adhesion (ITGA6 and ICAM3) and exosome and signaling molecules (GRB2 and PTPN6) (**Table S3, Tabs 5 and 6**) ^31^. The 12 upregulated transcripts in females were all mRNAs and were enriched for proteins involved with stress responses (DDX60L, RAD23B, USP15, HLA-C, and DNAJC7).

Finally, we assessed for each RNA X of each donor a Strong Translation Rate (STRx), (**Table S3 Tab 7**) ^31^, using:

STR_X_ = (Late Polysome) / (Free RNA + Monosome + Early Polysome). We identified 88 RNAs using these criteria (**Table S3 Tabs 8 and 9**) ^31^. Of these 88 RNAs, 81 were significantly upregulated in males and 7 were upregulated in females (**Table S3 Tabs 8 and 9**) ^31^. The 81 transcripts present in males included 3 pseudogenes, a novel transcript present within the sequence of phosphatidylinositol transfer protein membrane associated 2 (PITPNM2) and 76 mRNAs. These mRNAs were enriched for ribosome proteins, histone H4 lysine 20 (K20) demethylation, and respiratory burst (**Table S3, Tab 9**) ^31^. The 7 transcripts present in females included a mitochondrial pseudogene, and 6 mRNAs including myosin heavy chain 9 (MYH9). These data indicate that the TTRx, MTRx, and STRx groups in the male were enriched for transcripts associated with cell adhesion and signaling, whereas female neutrophils were modestly enriched for interferon signaling and stress responses.

### Male/female differences in RNAs only present in early and late polysomes

There were 523 transcripts that were only present in the polysome fractions, with 352 RNAs present in males and 314 present in females with infinite values for MTRx, indicating these RNA transcripts were only present in early and late polysomes and not in the free or monosome fractions (**Table S4, Tabs 1 and 2**) ^31^. Of the 523 transcripts, 209 were only detected in male neutrophils, 171 were only detected in female neutrophils, with 143 transcripts found in both male and female samples. Of the 352 transcripts in males, 14 were lncRNAs, 5 were mitochondrial tRNAs, 12 were processed pseudogenes, 3 encoded mitochondrial-derived humanin-like peptides, and 312 were protein coding transcripts of which 11 were encoded on the X chromosome (**Figure 6A and Table S4, Tab 3**) ^31^. The 312 protein-coding mRNAs were enriched for processes involved with cytoskeletal organization and cell motility, regulation of catabolic processes, and cellular differentiation (**Table S4, Tab 3)**. The 209 transcripts that were only detected in male neutrophils were also enriched for processes involved with cytoskeletal organization and cell motility. Of the 314 RNAs in females (**Table S4, Tab 4 and Figure 6B**) ^31^, 18 were lncRNAs, 4 were mitochondrial tRNAs, 12 were processed pseudogenes, 4 encoded mitochondrial-derived humanin-like peptides, and 277 were protein coding transcripts of which 6 were encoded on the X chromosome **(Table S4, Tab 4**). These transcripts were enriched for catabolic processes, regulation of RNA stability, and GTPase mediated signal transduction (**Table S4, Tab 4 and Figure 6B**). The 171 transcripts that were only detected in female neutrophils were also enriched for catabolic processes. For both male and female MTRx transcripts, there were no transcripts that were encoded on the Y chromosome.

**Figure 6.**
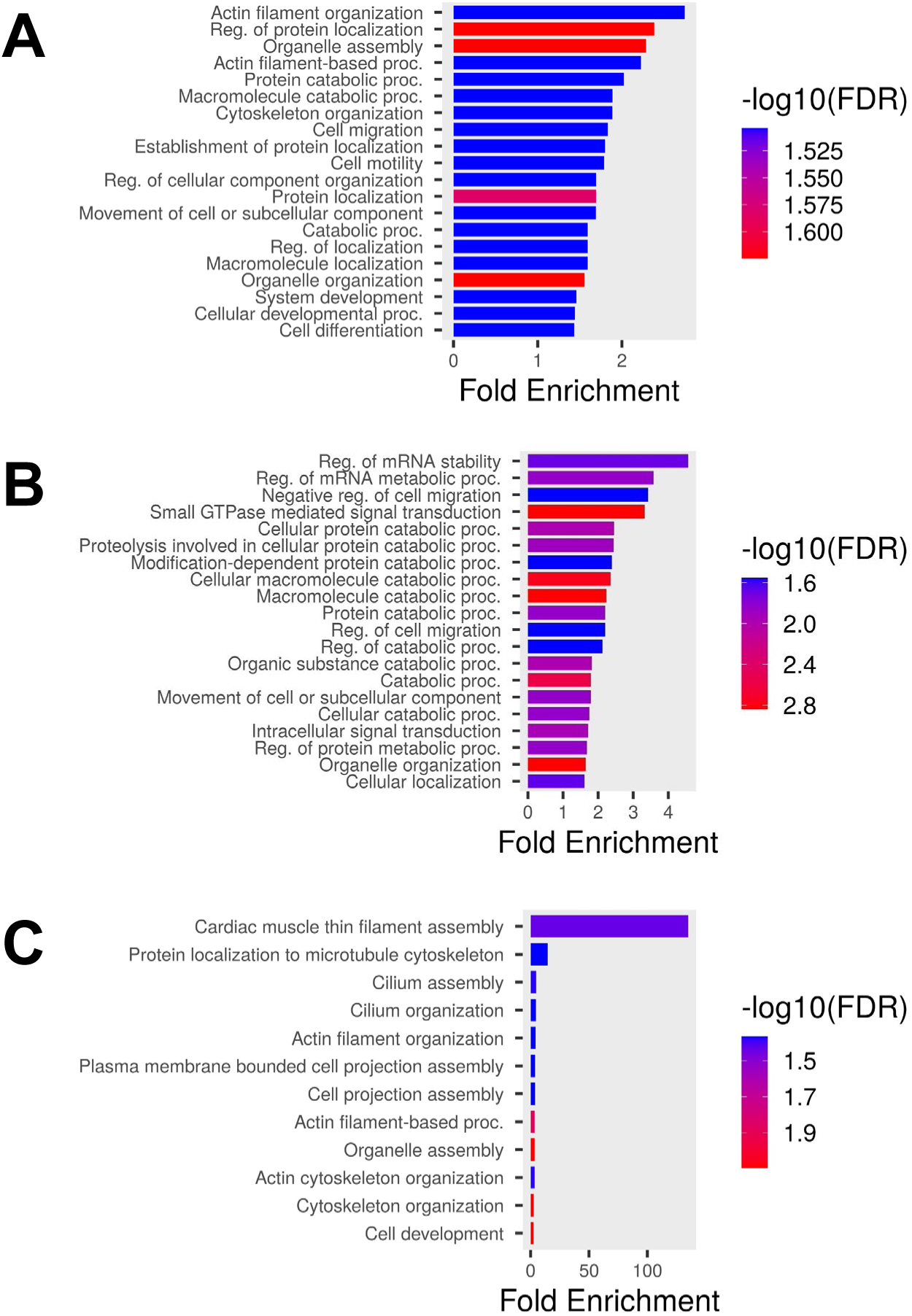
Analysis of RNA transcripts only present in polysomes. GO term analysis of enriched transcripts present only in the polysome fractions from **A)** male and **B)** female neutrophils with a moderate translational rate (MTRx). **C)** GO term analysis of enriched transcripts in male neutrophils with a strong translational rate (STRx).

### Male/female differences in RNAs only present in late polysomes

We also identified 128 RNAs in males and 52 in females with infinite values for STRx, indicating the RNA was only present in late polysomes and not in the free, monosome, or early polysome fractions (**Table S4, Tabs 5 and 6**) ^31^. Of the 128 STRx infinite value transcripts in males, 6 were lncRNAs, 1 was a mitochondrial tRNAs, 4 were processed pseudogenes, with 113 mRNAs of which 4 were encoded on the X chromosome (**Table S4, Tab 7**) ^31^. These transcripts were enriched for processes involved with cytoskeletal organization and cell motility (**Table S4, Tab 7 and Figure 6C**) ^31^. Of the 52 RNAs in females (**Table S4, Tab 8**) ^31^, 7 were lncRNAs, 3 were pseudogenes, and 41 were mRNAs of which 3 were encoded on the X chromosome (**Table S4, Tab 8**). These transcripts were weakly enriched for negative regulation of proteolysis and secretory vesicles (**Table S4, Tab8**). For both male and female transcripts only present in late polysomes, there were no transcripts that were encoded on the Y chromosome.

### Some transcripts were only present in male neutrophils

RNA isolated from whole neutrophils included 334 transcripts that were only detected in male neutrophils, compared to RNA isolated from female cells (**Table S1 Tab2 and 3**) ^31^ and 146 transcripts that were only detected in female neutrophils, compared to RNA isolated from whole male neutrophils **(Table S1 Tabs 2 and 4**) ^31^. We then determined if any of these 334 male transcripts were also detected in any of the 15,880 transcripts identified in any fraction of any of the 4 female samples from polysome fractionation **(Table S5 Tabs 5 and 7**) ^31^. Using these criteria, we identified 12 transcripts that were unique to male cells **(Table S5 Tab 8**) ^31^. Of these 12 transcripts, 2 were pseudogenes and 10 were mRNAs, with no transcripts that were encoded on the X or Y chromosome. The 10 mRNA transcripts (FMO4, UBXN8, KLF16, TMEM256, PIGC, WDR33, DEF8, TP53 [p53], LPCAT1, and TMEM243), included proteins that are present in the endoplasmic reticulum (UBXN8, PIGC, LPCAT1, FMO4, and TP53 [p53]).

From the male fractioned RNAs, there were 1,057 transcripts identified as being present in at least one fraction for all 3 donors **(Table S6 Tab 6**) ^31^. We then determined if any of these male transcripts were also detected in any female samples either from whole cells or polysome fractionation. Using these criteria, we identified 8 transcripts, with 1 lncRNA (ZNF407-AS1) and 7 mRNAs (CA8, TRIM21, ENOX1, CSNK2A2, XYLT2, CRNKL1, and STYL4), with SYTL4 encoded on the X chromosome **(Table S6 Tab 14**) ^31^. These 17 mRNA transcripts included RNA binding proteins (CRNKL1, WDR33, TP53, ENOX1, and TRIM21) and metal ion binding proteins (XYLT2, TP53, SYTL4, KLF16, TRIM21, DEF8, LPCAT1, and CA8), but were not enriched for any specific biological process.

### Some transcripts were only present in female neutrophils

For the 146 RNAs that were only present in whole female neutrophil RNA **(Table S1 Tab 4)**^31^, but not whole male neutrophil RNA, we determined if any of these female transcripts were also detected in any of the 15,046 transcripts identified in any fraction of any of the 3 male samples from polysome fractionation (**Table S5 Tab 6**) ^31^. Using these criteria, we identified 16 transcripts that were unique to female cells. Of these 16 transcripts, 2 were uncharacterized genes (TEC), 4 were lncRNAs, and 10 were mRNAs, with no transcripts that were encoded on the X or Y chromosome. The 10 mRNA transcripts (TSPOAP1, GATB, PCDHB4, DDX25, NT5DC3, TXNDC9, TMEM68, ADI1, DEFA3, and ADARB1), including proteins that regulate RNA (ADARB1, DDX25, GATB) (**Table S5 Tab 8**)^31^.

From the female fractioned RNAs, there were 376 transcripts identified as being present in at least one fraction for all 4 donors (**Table S6 Tab 12**) ^31^. None of these female transcripts were unique as these transcripts were present in either whole cell lysates or polysome fractionation of the male donors. Together, these data indicate that there are few sex-specific transcripts in male and female neutrophils.

## Discussion

Our data indicate that, as previously observed, ^15, 30^ male and female neutrophils have sex-based differences in levels of some RNAs isolated from whole cells. The donors were students recruited from the Texas A&M University population. We did not ask any donors if they were on any hormone treatment, so it is unclear if the observed differences are due to genetics or hormone treatment. Using a variety of different analysis schemes, we found that there were remarkably few transcripts that were unique to male or female neutrophils. Using the most rigorous scheme we observed 17 mRNA transcripts that were unique to male neutrophils, and 10 mRNA transcripts that were unique to all 4 female samples. Using alternative schemes, we observed that male neutrophils were enriched for transcripts encoding proteins involved in adhesion, leukocyte activation, ribosomes, and exosomes/vesicles. Female neutrophils were enriched for transcripts encoding proteins involved in mitochondrial function, cytokine production, and negative regulation of gene expression.

Polysome fractionation of RNAs has been used to determine translation efficiency in many cell types, but this is rare for neutrophils ^33^. In male neutrophils, we observed that RNAs not associated with a ribosome (the free fraction) were enriched for RNAs encoding mitochondrial respiratory chain complex proteins, suggesting that in male neutrophils mitochondrial processes are not actively maintained. The free RNA fraction in female cells was enriched for RNA processing and regulation RNAs, suggesting that translation is a less active process in female cells.

Analysis of RNAs enriched in the polysomes or only present in polysomes, and thus likely to be involved with active translation, indicated that male neutrophils have active translation of RNAs associated with cytoskeletal organization and cell motility, whereas female neutrophils actively translate mRNAs associated with interferon signaling and metabolic processes such as catabolism. In addition, we observed that the polysome fractions of female neutrophils were enriched for RNAs encoding proteins associated with mitochondria, whereas male neutrophils had an enrichment of mitochondrial RNAs in the free RNA fraction. Analysis of the TTRx, MTRx, and STRx data, which determine translation efficiency, indicated that male neutrophils had higher translation rates of transcripts encoding proteins involved with cell adhesion and signaling, whereas female neutrophils had higher translation rates of transcripts encoding proteins involved with interferon signaling and stress responses. These data suggest a clear difference in the biology of male and female neutrophils and may also help explain the differences in the responses of male and female neutrophils to infection ^5, 6, 43, 44^.

For some cell types, RNA transcript levels correlate with protein levels ^45^, but for many cell types including neutrophils, specific RNA transcript levels do not match the corresponding protein levels ^46, 47^. Discrepancies between RNA transcript levels and protein abundance may be attributed to the association of RNA transcripts with polysomes ^38, 48^. Cells with high levels of RNA transcripts that are not associated with ribosomes (free RNA) or transcripts only associated with monosomes will not be effectively translated into proteins, whereas low copy number RNA transcripts associated with polysomes are likely to generate many protein copies ^49^. Protein levels are also dependent on protein turnover and degradation rates ^45–47, 50^. There are also sex differences in protein degradation, which may also influence protein levels in males and females^51, 52^. We have previously shown that neutrophils from males have higher levels of proteins that regulate translation and signaling cascades, whereas neutrophils from females have higher levels of proteins that regulate metabolic processes and proteolytic pathways ^19^. Although we did not identify RNA transcripts that encode proteins we had previously identified to be elevated in male or female neutrophils ^19^, we did find that male neutrophils were enriched for transcripts encoding proteins involved in adhesion and leukocyte activation, and female neutrophils were enriched for transcripts encoding proteins involved in mitochondrial function and negative regulation of gene expression. These data suggest that although RNA transcripts and proteomics data may not match, grouping of datasets into modules or clusters ^26, 28, 46^ may reveal similarities between RNA and protein datasets.

We previously observed that male neutrophils have a more rapid response to chemorepellents ^22, 53^. We found that male neutrophils also had increases in many RNAs associated with leukocyte adhesion and chemotaxis, suggesting that in male cells the faster response may be due to increased translation of RNAs associated with these processes. These differences may also contribute to the observed sex-based differences in neutrophil behavior and neutrophil-associated disease incidence and severity, and suggests that therapies that affect neutrophil biology may need to be modified for male or female patients ^54–57^.

## Declarations

### Ethics approval and consent to participate

The studies involving human participants were reviewed and approved by the Texas A&M University Institutional Review Board (IRB #2017-0792D). Written informed consent to provide blood for this study was provided by the donor.

### Consent for publication

Not applicable.

## Availability of data and materials

The datasets used and/or analyzed during the current study are available from the corresponding author on reasonable request. RNA-seq data were uploaded to GEO (accession number GSE288976).

## Authors’ Contributions

K.M.C. and D.P. designed, performed, analyzed experiments, and wrote the paper. S.A.K. performed experiments. R.H.G. designed experiments and wrote the paper.

## Acknowledgements

We thank the volunteers who donated blood to perform these experiments, the phlebotomy staff at the Texas A&M Beutel Student Health Center, the Texas A&M Institute for Genome Sciences and Society (TIGSS) Experimental Genomics Core for RNA library sequencing. We also thank Sumeen Gill, Md Salman Zahir Uddin and Shivani Ram Gianani for helpful comments on the manuscript.

## Author Information

Darrell Pilling, Kristen M. Consalvo, Sara A. Kirolos, and Richard H. Gomer Department of Biology, Texas A&M University, College Station, TX 77843-3474 USA

---

**Figure S1.**
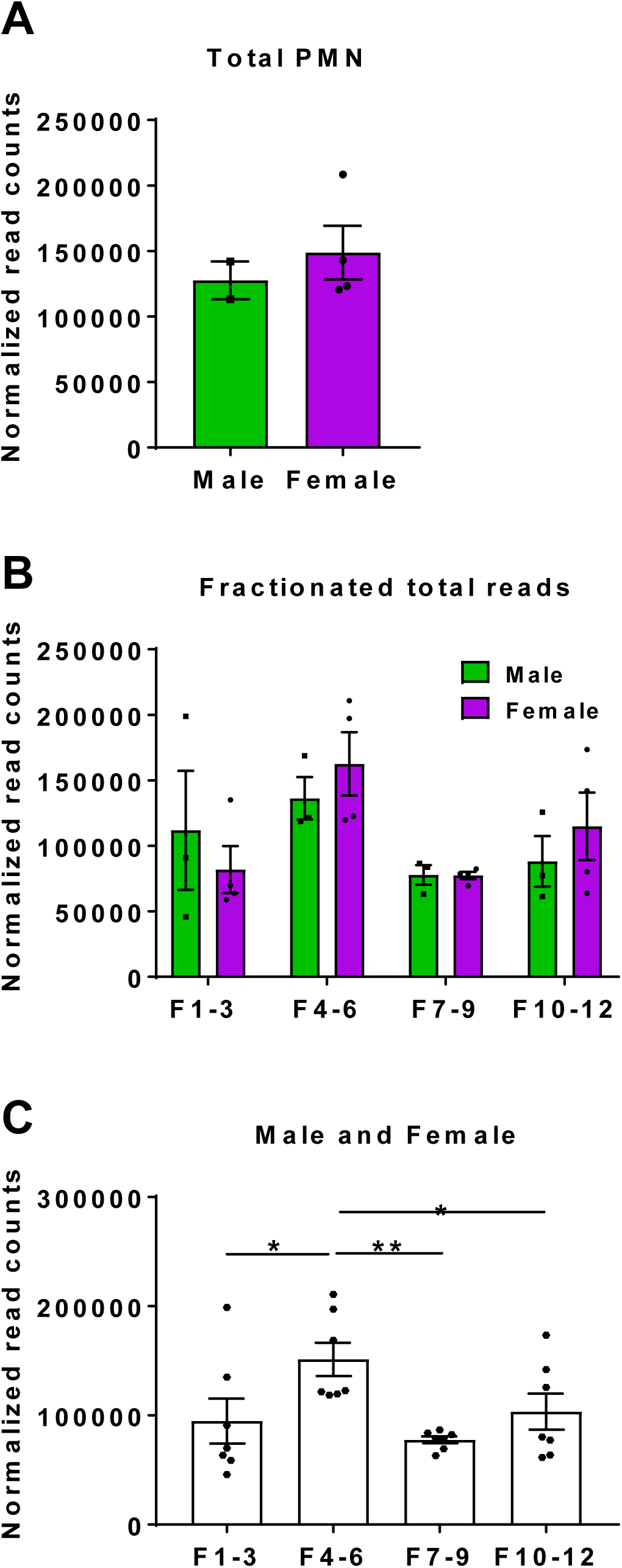
Analysis of read counts from male and female neutrophils. **A)** Number of normalized read counts from total RNA isolated from male and female neutrophils (n = 2 male and 4 female donors). **B)** Number of normalized read counts from RNA isolated from polysome fractionation of male and female neutrophils. **C)** Combined normalized read counts from RNA isolated from polysome fractionation of male and female neutrophils (n = 3 male and 4 female donors). Bars and error bars show mean ± SEM. * indicates p < 0.05 and ** indicates p < 0.01 (1-way ANOVA with Tukey’s test).

**Table S1: Analysis of RNA-seq from male and female neutrophils. Tab 1)** RNA-seq of total RNA. M2 and M3 are male donors, F1-F4 are female donors. Values are read counts from RNA-seq of total mRNA of 2,175 transcripts that had detectable RNA sequences (reads) in either both male donors or in at least 2 of 4 female donors. **Tab 2)** RNA-seq values ranked by total abundance of RNAs from combined male and female samples. **Tab 3)** Analysis of transcripts only identified in male neutrophils. **Tab 4)** Analysis of transcripts only identified in female neutrophils. **Tab 5)** Data sets used for volcano plots in Figure 1B.

**Table S2: Analysis of sex-based differences in RNA transcripts from free RNA, monosome, early and late polysome fractions of male and female neutrophils.** Values are normalized read counts from RNA-seq analysis of 3,056 transcripts that had at least 3 detectable RNA sequences (reads) in at least one fraction either both male donors or in at least 2 of 4 female donors. **Tab 1)** Analysis of RNAs identified from free RNA, monosome, and early and late polysome fractions of neutrophils from 3 male (M1-M3) and 4 female (F1-F4) donors. **Tab 2)** Identification of transcripts only present in male or female samples. **Tab 3)** Mean values from **Tab 2** for ClustVis analysis. **Tab 4)** Processed data from ClustVis analysis for heatmap (Figure 4). **Tab 5)** Analysis of >2 fold changed transcripts from ClustVis. **Tab 6)** Transcripts ranked above 1.0 from Clustvis. **Tab 7)** Transcripts ranked below 1.0 from Clustvis.

**Table S3: Analysis of sex-based differences in RNA translation rates.** Values are normalized read counts from RNA-seq analysis of 3,056 transcripts that had at least 3 detectable RNA sequences (reads) in at least one fraction either both male donors or in at least 2 of 4 female donors. **Tab 1)** For each donor, to calculate the total translation rate (TTRx) the normalized read counts from each individual donor of monosomes, early polysomes, and late polysome fractions were combined and then the values were divided by the reads from each donors’ free RNA fraction. **Tab 2)** List of those transcripts that were significantly different between male and female neutrophils. **Tab 3)** Ranked list from **Tab 2** of transcripts that were significantly different between male and female neutrophils. **Tab 4)** For each donor, to calculate the moderate translation rate (MTRx) the normalized read counts from each individual donor of early and late polysome fractions were combined and then the values were divided by the combined reads from each donors’ free RNA and monosome fraction. **Tab 5)** List of the MTRx transcripts that were significantly different between male and female neutrophils. **Tab 6)** Ranked list from **Tab 5** of MTRx transcripts that were significantly different between male and female neutrophils. **Tab 7)** For each donor, to calculate the strong translation rate (STRx) the normalized read counts from each individual donor late polysome fractions were combined and then the values were divided by the combined reads from each donors’ free RNA, monosome, and early polysome fraction. **Tab 8)** List of the STRx transcripts that were significantly different between male and female neutrophils. **Tab 9)** Ranked list from **Tab 8** of STRx transcripts that were significantly different between male and female neutrophils.

**Table S4: Analysis of sex-based differences in RNA transcripts only present in polysomes.** Analysis of the transcription rates from **Table S3** indicated there were many transcripts that were only present in the polysome fractions. These data gave infinite values for the MTRx and STRx calculations, that is there were no transcript detectable RNA sequences (reads) in the free RNA or monosome fractions but there were transcript reads in the early and late polysome fractions. **Tab 1)** Calculations to identify for each donor, those RNAs where the polysome fractions were divided by zero (infinity values), as in no reads in the free RNA or monosome fractions. **Tab 2)** Ranked list from **Tab 1** of transcripts that had infinity values for the moderate translation rate (MTRx). **Tab 3)** Analysis of male transcripts from **Tab 2** with infinity values for the MTRx. **Tab 4)** Analysis of female transcripts from **Tab 2** with infinity values for the MTRx. **Tab 5)** Calculations to identify for each donor, those RNAs where the late polysome fractions were divided by zero (infinity values), as in no reads in the free RNA, or monosome, or early polysome fractions. **Tab 6)** Ranked list from **Tab 5** of transcripts that had infinity values for the strong translation rate (STRx). **Tab 7)** Analysis of male transcripts from **Tabs 5 and 6** with infinity values for the STRx. **Tab 8)** Analysis of female transcripts from **Tabs 5 and 6** with infinity values for the STRx. **Tab 9)** Venn diagrams analysis of the overlap of transcripts in male and female MTRx and STRx datasets.

**Table S5: Analysis of sex-based differences in RNA transcripts from whole cell and polysome fractionated samples. Tab 1)** RNA-seq of total RNA. M2 and M3 are male donors, F1-F4 are female donors. **Tab 2)** Ranked data of all 9,344 transcripts with at least one sequence read. **Tab 3)** All 1301 transcripts present in both male total lysate samples. **Tab 4)** All transcripts present in n=2, n=3, or n=4 female total lysate samples. **Tab 5)** All transcript reads from polysome fractionated male and female samples. **Tab 6)** Ranked male transcripts. **Tab 7)** Ranked female transcripts. **Tab 8)** Identification of male only transcripts that were absent from female whole cell lysates or fractionated lysates, and identification of female only transcripts that were absent from male whole cell lysates or fractionated lysates.

**Table S6: Analysis of sex-based differences in RNA transcripts from polysome fractionated samples. Tab 1)** Ranked datasets of all 5,241 transcripts with at least 3 reads present in any fraction of any male sample. **Tab 2)** Ranked datasets of all 754 transcripts with 3 transcript reads present in the free RNA (F1-3) of male samples. **Tab 3)** Ranked datasets of all 616 transcripts with 3 transcript reads present in the monosome fraction (F 4-6) male samples. **Tab 4)** Ranked datasets of all 984 transcripts with 3 transcript reads present in the early polysome fraction (F 7-9) male samples. **Tab 5)** Ranked datasets of all 829 transcripts with 3 transcript reads present in the late polysome fraction (F 10-12) male samples. **Tab 6)** List and Venn diagram of transcripts present in at least one fraction for all 3 male donors. **Tab 7)** Ranked datasets of all 5,892 transcripts with at least 3 reads present in any fraction of any female samples. **Tab 8)** Ranked datasets of all 754 transcripts with 3 transcript reads present in the free RNA (F1-3) sample. **Tab 9)** Ranked datasets of all 939 transcripts with 3 transcript reads present in the monosome fraction (F 4-6) of female samples. **Tab 10)** Ranked datasets of all 1,828 transcripts with 3 transcript reads present in the early polysome fraction (F 7-9) of female samples. **Tab 11)** Ranked datasets of all 1,155 transcripts with 3 transcript reads present in the late polysome fraction (F 10-12) of female samples. **Tab 12)** List and Venn diagram of transcripts present in at least one fraction for all 4 female donors. **Tab 13)** List and Venn diagram of transcripts present in at least one fraction for at least 3 of the 4 female donors. **Tab 14)** List of transcripts found only in male and female samples.

